# Gut Microbiome Biomarkers in Adolescent Obesity: a Regional Study

**DOI:** 10.1101/745489

**Authors:** Xuefeng Gao, Binbin Wu, Yonglong Pan, Shaoming Zhou, Ming Zhang, Yunpeng Cai, Yan Liang

## Abstract

**Purpose:** This study aimed to characterize the gut microbiota in obese Shenzhen adolescents, and evaluate the influence of gender on BMI-related differences in the gut microbiome.

**Methods:** Physical examinations, blood pressure measurement, serological assay, and body composition evaluation were conducted on two-hundred and five adolescents from Shenzhen. Fecal microbiome composition was profiled via 16S rRNA gene sequencing. A Random Forest (RF) classifier model was built to distinguish the BMI categories based on the gut bacterial composition.

**Results:** Fifty-six taxa consisting mainly of *Firmicutes* were identified that having significant associations with BMI; two OTUs belonging to *Ruminococcaceae* and one belonging to *Lachnospiraceae* had relatively strong positive correlations with body fate rate, waistline, and most of serum biochemical parameters. Based on the 56 BMI-associated OTUs, the RF model showed a robust classification accuracy (AUC 0.96) for predicting the obese phenotype. Gender-specific differences in the gut microbiome composition was obtained, and a lower relative abundance of *Odoribacter* was particularly found in obese boys. Functional analysis revealed a deficiency in bacterial gene contents related to PPAR signaling pathway in obese subjects for both genders; significantly lower levels of adipocytokine signaling pathway and ethylbenzene degradation were particularly detected in obese girls.

**Conclusions:** This study revealed unique features of gut microbiome in terms of microbial composition and metabolic functions in obese Shenzhen adolescents. The effect of geographical location, age and gender on the gut microbiome should be carefully considered in case–control studies.

## INTRODUCTION

Obesity during childhood and adolescence is associated with cardiovascular disease and metabolic syndrome later in life, and has become a significant public health concern worldwide (Collaboration NCDRF. 2017). The human gut microbiome is being increasingly recognized as an important factor in the development of obesity (Ley et al. 2006; Shen et al. 2013), and it is highly involved in the host calorie harvest and energy homeostasis (Rios-Covian et al. 2016). Alterations in the gut microbiota caused by such as antibiotics exposure, is linked to a variety of metabolic diseases including obesity, type 1 and type 2 diabetes (Cox and Blaser. 2015). Studies from both animal and human have suggested divergences in the gut microbiome composition result in different weight gain post same diet (Le Chatelier et al. 2013; Thaiss et al. 2016). In addition, some gut microbes may stimulate chronic low-grade inflammation by such as producing lipopolysaccharides, thereby contributing to obesity and insulin resistance (Boulange et al. 2016). Moreover, modulation of the gut microbiota by fiber supplementation or fecal microbiota transplantation can suppress inflammation and improve insulin sensitivity, suggesting the vital role of the gut microbiota in etiology of metabolic syndrome (Cani et al. 2008).

Investigations have been performed to identify the gut microbiota markers of obesity; However, no consistent pattern was not obtained from different studies. For example, both Bäckhed et al. (Backhed et al. 2004) and Turnbaugh et al (Turnbaugh et al. 2006). found that a decreased ratio of Bacteroidetes to Firmicutes in people with obesity, but failed to be supported by the other human studies (Shi et al. 2006; Duncan et al. 2008). Indeed, the gut microbiome composition appears to be shaped by host genetics, age, gender, geographical locations, and other environmental factors. By characterizing the gut microbiota of 7,009 individuals from 14 districts within the Guangdong province of China, He et al. showed that host location is the strongest explanatory factor to the microbiota variations (He et al. 2018). Our previous study demonstrated BMI differences in the gut microbiota are gender specific (Gao et al. 2018). Thus, the region, age and gender should be considered when choosing controls to compare the gut microbiome of disease cases.

By accounting for the geographical location and age as confounding factors, this study characterizes the composition and functions of the gut microbiota in obese Shenzhen adolescents, and evaluated the influence of gender on the BMI-related differences in the gut microbiome.

## MATERIAL AND METHODS

### Study cohort

The study began following approval from the Institutional Review Board of the Shenzhen Institutes of Advanced Technology, Chinese Academy of Sciences (SIAT-IRB-131115-H0032), and was registered with ClinicalTrials.gov (number NCT02539836). All individuals participating in the study received informed consent from their guardians. Two hundred and seventeen Chinese adolescents were recruited at Shenzhen Children’s Hospital, between September and December 2015. Written informed consent was obtained from parents of the subjects before participation. Subjects with any of the criteria below were excluded from the present study:

1. Type 1 or type 2 diabetes.
2. Antibiotic use in the two months prior to sampling.
3. Long-term use of medication (e.g. antihypertensive drugs).
4. Diarrhea.
5. Long-term constipation.

According to the WHO BMI-for-age percentile growth charts (growth reference 5-19 years; https://www.who.int/growthref/who2007_bmi_for_age/en/), we classified each participant into normal weight (N), overweight (OW), or obese (OB). General characteristics of the subjects, body composition, and serological test results for each BMI group are shown in Table 1.

**Table 1.** Characteristics of the study population.

| Characteristics | Normal | Over Wight | Obesity | p-Value (ANOVA) |
| --- | --- | --- | --- | --- |
| Number (F/M) | 65 (30/35) | 53(19/34) | 87 (19/68) |  |
| Age (years) | 13.91± 1.61 | 13.26 ± 1.50 | 12.90 ± 1.41 | 2.75E-04 |
| Age Range (years) | 11–16 | 11–16 | 11–16 |  |
| **Anthropometric** |  |  |  |  |
| Weight (kg) | 50.60 ± 9.67 | 62.88 ± 8.85 | 74.62 ± 11.50 | 2.99E-31 |
| Height (m) | 163.03 ± 9.31 | 162.81 ± 8.94 | 163.92 ± 8.79 | 7.34E-01 |
| BMI | 18.87 ± 1.96 | 23.60 ± 1.42 | 27.65 ± 2.53 | 1.78E-63 |
| Waistline (cm) | 66.91 ± 10.72 | 78.19 ± 6.81 | 88.36 ± 7.16 | 9.40E-45 |
| **Blood Pressure** |  |  |  |  |
| Systolic blood pressure (mm Hg) | 114.28 ± 14.48 | 121.87 ± 11.93 | 125.76 ± 12.48 | 1.20E-06 |
| Diastolic blood pressure (mm Hg) | 70.78 ± 8.17 | 72.79 ± 10.24 | 76.59 ± 8.74 | 3.84E-04 |
| **Body composition** |  |  |  |  |
| Body fat | 7.67 ± 4.12 | 14.91 ± 4.12 | 21.81 ± 10.60 | 8.69E-40 |
| Body fat rate | 0.15 ± 0.07 | 0.23 ± 0.08 | 0.29 ± 0.13 | 6.94E-34 |
| Body muscle | 40.67 ± 13.85 | 45.08 ± 14.78 | 49.55 ± 22.69 | 2.95E-09 |
| Body muscle rate | 0.8 ± 0.24 | 0.71 ± 0.21 | 0.66 ± 0.29 | 8.49E-22 |
| Body mineral salts | 3.04 ± 1.05 | 3.66 ± 1.18 | 4.32 ± 2.01 | 1.62E-03 |
| Body mineral salts rate | 0.06 ± 0.02 | 0.06 ± 0.02 | 0.06 ± 0.03 | 8.19E-34 |
| Body water | 31.86 ± 10.87 | 35.10 ± 11.48 | 38.78 ± 17.99 | 4.09E-09 |
| Body water rate | 0.62 ± 0.19 | 0.55 ± 0.17 | 0.51 ± 0.23 | 1.69E-27 |
| Body protein | 9.28 ± 3.17 | 9.98 ± 3.30 | 10.76 ± 5.00 | 1.62E-05 |
| Body protein rate | 0.18 ± 0.06 | 0.16 ± 0.05 | 0.14 ± 0.06 | 2.01E-31 |
| **Serological investigations** |  |  |  |  |
| ALB | 47.62 ± 4.33 | 46.86 ± 3.99 | 47.69 ± 4.46 | 5.06E-01 |
| ALT | 13.55 ± 8.60 | 17.06 ± 12.79 | 27.66 ± 22.58 | 1.26E-06 |
| ApoA1 | 1.38 ± 0.17 | 1.31 ± 0.17 | 1.33 ± 0.17 | 9.49E-02 |
| ApoB | 0.60 ± 0.15 | 0.64 ± 0.16 | 0.7 ± 0.16 | 1.91E-04 |
| BUN | 3.85 ± 1.07 | 3.95 ± 0.86 | 4.08 ± 0.83 | 3.03E-01 |
| CER/CP | 24.35 ± 4.64 | 28.07 ± 5.60 | 29.06 ± 6.39 | 3.39E-06 |
| CK | 123.83 ± 59.79 | 133.43 ± 68.69 | 179.41 ± 184.46 | 2.00E-02 |
| CK-MB | 1.52 ± 0.67 | 1.60 ± 0.80 | 1.90 ± 1.13 | 3.35E-02 |
| C-peptide | 1.88 ± 1.13 | 2.48 ± 1.80 | 2.82 ± 1.93 | 3.47E-03 |
| Cr | 55.93 ± 13.71 | 54.46 ± 11.00 | 54.08 ± 11.47 | 6.32E-01 |
| CRP | 0.46 ± 0.55 | 1.39 ± 1.91 | 2.23 ± 3.16 | 2.94E-05 |
| Hb | 134.95 ± 18.06 | 137.30 ± 11.03 | 136.94 ± 14.66 | 6.35E-01 |
| HDL | 1.22 ± 0.23 | 1.12 ± 0.23 | 1.09± 0.21 | 1.18E-03 |
| INS | 10.35 ± 8.00 | 16.28 ± 15.91 | 22.04 ± 17.67 | 1.56E-05 |
| LDL | 2.34 ± 0.56 | 2.55 ± 0.68 | 2.69 ± 0.55 | 2.15E-03 |
| PLC | 291.12 ± 64.73 | 312.93 ± 57.37 | 329.56 ± 63.03 | 1.04E-03 |
| PUFA | 0.45 ± 0.20 | 0.50 ± 0.21 | 0.57± 0.27 | 6.06E-03 |
| RBC | 4.91 ± 0.53 | 4.90 ± 0.36 | 5.02 ± 0.42 | 1.94E-01 |
| TBIL | 9.85 ± 4.52 | 8.72 ± 4.92 | 7.23 ± 3.11 | 5.64E-04 |
| TC | 3.93 ± 0.83 | 4.07 ± 0.85 | 4.37 ± 0.77 | 3.47E-03 |
| TG | 1.09 ± 0.60 | 1.59 ± 1.17 | 1.76 ± 0.94 | 1.02E-03 |
| TP | 76.90 ± 7.98 | 75.70 ± 6.00 | 76.41 ± 5.50 | 6.11E-01 |
| UA | 337.71 ± 87.60 | 398.79 ± 120.42 | 425.78 ± 102.84 | 2.55E-06 |
| WBC | 7.61 ± 1.76 | 8.60 ± 2.05 | 8.70 ± 1.67 | 6.79E-04 |

Values represent mean ± standard error of the mean.

### Sample collection, DNA extraction, and 16S rRNA gene Sequencing

Venous blood samples were collected at Shenzhen Children’s Hospital upon annual physical examination. Serological assays were performed to determine white blood cell (WBC), red blood cell (RBC), Hemoglobin (Hb), proinsulin-like component (PLC), alanine aminotransferase (ALT), total bilirubin (TBIL), total protein (TP), albumin (ALB), Cerutoplasmin (CER/CP), ceruloplasmin (CRP), blood urea nitrogen (BUN), creatinine (Cr), urine acid (UA), creatine kinase (CK), creatine kinase-myoglobin (CK-MB), insulin (INS), C-peptide, triglycerides (TG), total cholesterol (TC), high-density lipoprotein (HDL), low-density lipoprotein (LDL), apolipoprotein A1 (Apo-A1), apolipoprotein B (Apo-B), and polyunsaturated fatty acid (PUFA).

Each participant donated ∼10g of fresh stool and placed inside a sterile plastic bag with ice pack. These samples were homogenized to a uniform consistency, and DNA was routinely extracted from 0.3g fecal material using TIANamp Stool DNA Kit (TIANGEN BIOTECH, cat. #DP328-02, Beijing, China), following the manufacturer’s instructions. DNA was quantified using a dsDNA HS assay on a Qubit 3.0 (Thermo Fisher Scientific, USA). The universal primers (forward: 5′-AYTGGGYDTAAAGNG-3′, reverse: 5′-TACNVGGGTATCTAATCC-3′) were used to PCR amplify the isolated genomic DNA for the V3–V4 16S rDNA hypervariable regions. The PCR products were sequenced by an Illumina MiSeq (Illumina, Inc, San Diego, CA) using the 2x300 bp paired-end protocol.

### 16S rRNA Gene Sequencing Analysis

The raw sequencing data was quality filtered and demultiplexed using QIIME 1.9.1 (Caporaso et al. 2010). Chimeric sequences were identified and removed using UCHIME (Edgar et al. 2011). Two-hundred and five samples which passed the quality control were incorporated into analysis. Operational taxonomic units (OTUs) were clustered using a closed-reference picking protocol with the UCLUST algorithm based on 97% nucleotide similarity. Microbial OTUs were annotated with the Greengenes reference database 13_8. OTU counts were processed with total sum normalization (TSS) followed by cumulative-sum scaling (CSS) by using Calypso (Zakrzewski et al. 2017). The relative abundances were log2 transformed to account for the non-normality of taxonomic counts data.

The biodiversity was measured by number of OTUs, Chao1 index, Shannon index, and Inverse Simpson index. We performed Principal Coordinate Analysis (PCoA) to determine whether the samples could be separated on the basis of BMI and gender. Significance of composition difference among groups was determined using analysis of variance (ANOVA) followed by Wilcoxon test for pairwise comparisons. We used Spearman correlations to identify the BMI-associated taxa. We considered OTUs appeared in more than 25% of the samples, that had significant correlation (p<0.01) with BMI. PICRUSt (Phylogenetic Investigation of Communities by Reconstruction of Unobserved States) (Langille et al. 2013) was applied to obtain the KEGG (Kyoto Encyclopedia of Genes and Genomes) pathways through predicting metagenome content from 16S rRNA gene surveys.

A Random Forest (RF) model was constructed by using R-package randomForest (Liaw and Wiener. 2002) to perform supervised classification of the three BMI categories. Half of the dataset was used for building and training the RF model, and the other half for testing. The optimal number of variables (*mtry*) randomly sampled as candidates at each split was accessed by *tuneRF* function. We calculated the interpolated area under the ROC curve (area under the receiver operating characteristic curve, AUC) for each classifier based on the cross-validation testing results.

## RESULTS

### BMI Is Associated with Compositional Changes in The Gut Microbiome of Adolescents

After quality filtering, a total of 17,323,193 sequencing reads were obtained from 205 fecal samples. Taxa that have less than 0.01% relative abundance across all samples were excluded. An overall of 518 OTUs were identified, which were grouped in 9 phylum and 69 genera. Richness (number of OTUs and Chao1 indexes) and alpha diversity (Shannon and Inverse Simpson indexes) were estimated at the OTU level with samples rarefied to depth of 37,027 reads (the lowest number of sequences). The richness of was higher in the overweight subjects compared with those with normal weight (Figure S1A, B). No significance in alpha diversity was obtained among the three BMI groups (Figure S1C, D). With respect to beta diversity, PCoA of Bray-Curtis dissimilarity index demonstrated that the overall gut microbiome composition was not able to be stratified by BMI (Figure S2).

Differential taxa abundance was analyzed using one-way ANOVA. At the phylum level, *Verrucomicrobia* was more abundant in obese adolescents (Figure 1A). At the family level, obese adolescents had increased abundances of *Actinomycetaceae* (including genera *Actinobacteria* and *Collinsella*; Figure 1C), *Clostridiaceae* (including genera *Clostridium* and *SMB53*; Figure 1C), *Cytophagaceae* (including *Rhodococcus* genus; Figure 1C), *Streptococcaceae* (including *Streptococcus* genus; Figure 1C), and *Veillonellaceae*, and decreased abundances of *Christensenellaceae* and *Rikenellaceae* (Figure 1B). The relative abundance of *Nocardiaceae* family was particular higher in the overweight subjects. In addition, *Erwinia* belonging to *Enterobacteriaceae* family was found with a higher relative abundance in obese adolescents (Figure 1C).

**Figure 1.**
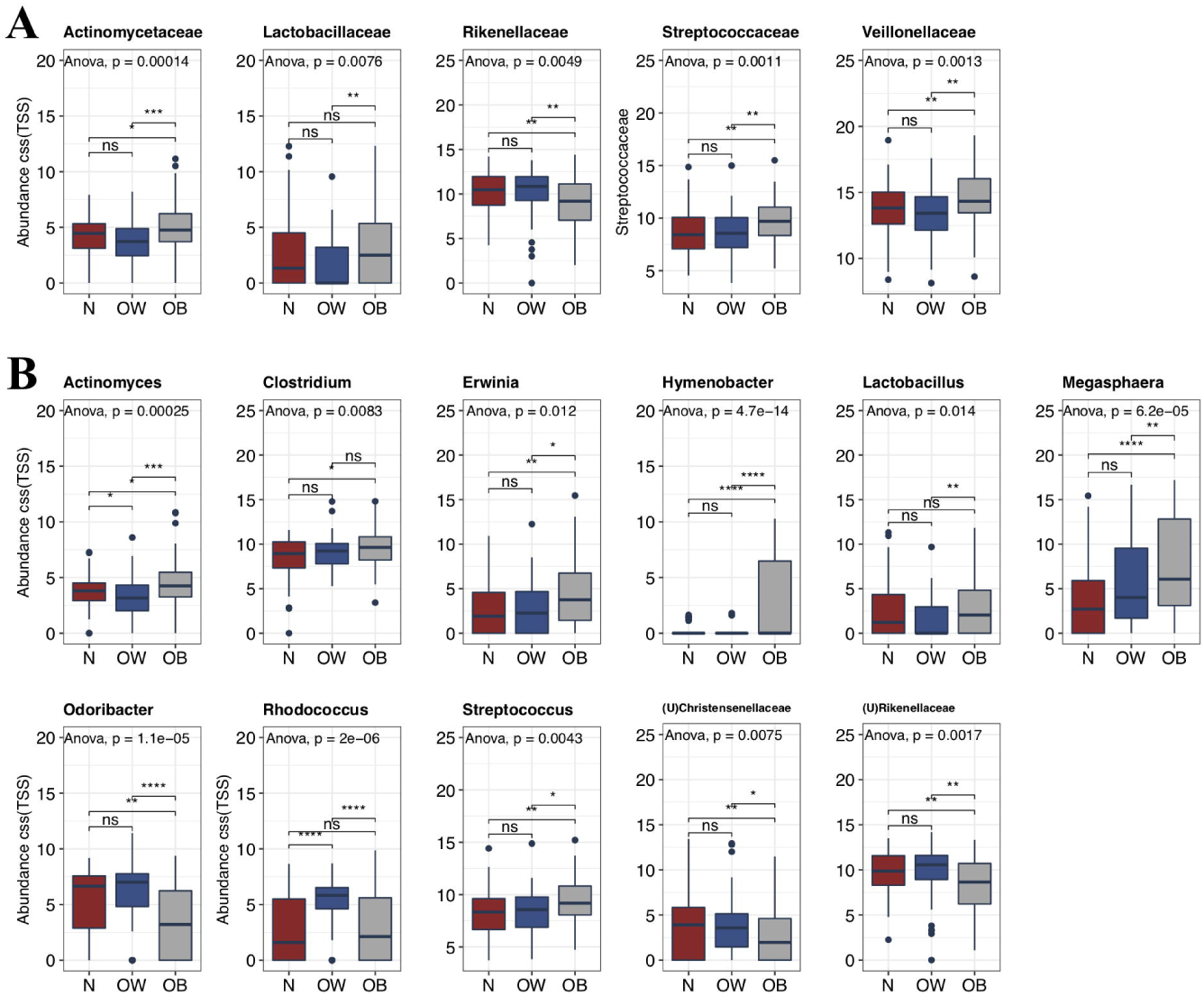
Differences in fecal microbiome constituents of normal weight, overweight, and obese Shenzhen adolescents. Bacterial taxa at the **(A)** family and **(B)** genus levels demonstrated significantly different abundances across the three BMI groups. Each bar plot indicates the mean proportion of sequences assigned to a feature in each group. Whiskers represent 1.5* inter-quartile range. Relative abundances were analyzed by one-way ANOVA followed by Wilcoxon test for pairwise comparisons. **P < 0.01 and ***P < 0.001. N, normal weight; OW, overweight; OB, obese.

Fifty-six taxa were identified with significant correlations with BMI with most of the members belonging to the *Firmicutes* phylum (48 OTUs), and 17 and 12 taxa part of the *Ruminococcaceae* and *Lachnospiraceae* families, respectively (Table S1, S2, S3). For each content of serological surveys and boy composition, we used Spearman correlation method to identify their associations with the 56 BMI-associated taxa. Three Firmicutes taxa (two OTUs belonging to *Ruminococcaceae* and one belonging to *Lachnospiraceae*) were found having relatively strong positive correlations with BMI, body fate rate, waistline, and inverse correlations with body water, protein, muscle, and mineral salts rates (Figure 2A). Notably, these three taxa also positively correlated with most of serum biochemical parameters, except TBIL. Notably, a few of *Ruminococcaceae* bacteria were inversely correlated with BMI, including *Faecalibacterium prausnitzii*.

**Figure 2.**
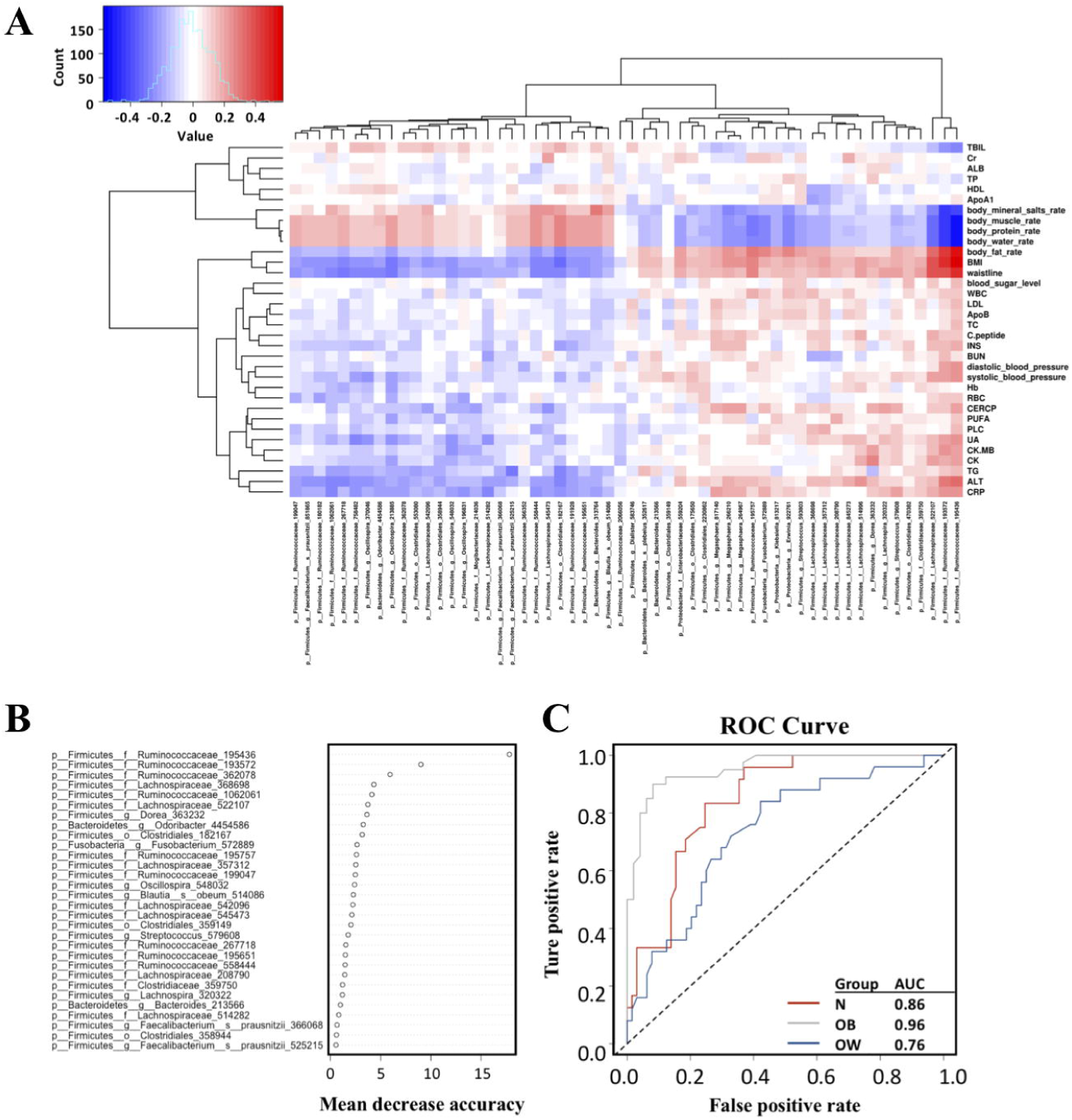
Identifying BMI-associated bacterial taxonomic biomarkers Shenzhen adolescents. **(A)** Heatmap of Spearman correlations between 54 BMI-associated bacterial taxa and serum biochemical parameters and body composition. **(B)** Thirty BMI-discriminatory bacterial taxa were identified by the RF model which were listed in rank order of their contribution to the classification accuracy (mean decrease accuracy). **(C)** ROC curve of the RF model.

To further investigate the BMI-associated bacterial taxonomic biomarkers in the gut microbiome in Chinese adolescents, we built a RF model to classify the phenotypes based on the 56 OTUs identified by the Spearman correlation. Tuning of the RF model resulted in *mtry*=14 for *ntree*=500. The 30 most BMI-discriminatory taxa identified by the RF model were shown in Figure 2B in rank order of their contribution to the predictive accuracy. We calculated the interpolated area under the receiver operating characteristic (ROC) curves (AUC) for the classifier based on the cross-validation testing results. We successfully classified obese adolescents with a high classifiability (Figure 2C; AUC=0.96). Adolescents with normal weight and those overweight were also able to be classified, with AUC of 0.86 and 0.76 for subjects with normal weight and overweight, respectively.

### Gut Microbiota Gene Content Associated with PPAR Signaling and Adipocytokine Signaling Are Reduced in Obese Adolescents

We also used Spearman correlations to identify the BMI-associated functional profile of microbial community 16S rRNA sequence data by using PICRUSt (Langille et al. 2013). We selected seven KEGG pathways that had significant correlation (p<0.01) with BMI (Table S4, S5, S6). For each of the KEGG pathways that significantly associated with BMI, we further used Spearman correlations to identify their associations with the serological parameters and body composition (Figure 3A). The predicted gene content of related to peroxisome, PPAR signaling pathway, and adipocytokine signaling pathway showed inverse associations with BMI, body fat rate, and waistline, and positive correlation with body water rate, protein rate, muscle rate, and mineral salts rate. In particular, PPAR signaling pathway and adipocytokine signaling pathway were found significantly downregulated in the obese adolescents comparing to the normal weight and overweight counterparts (Figure 3B). Additionally, these pathways were inversely correlated with CER/CP, blood sugar level, (both systolic and diastolic) blood pressure, CRP, WBC, PUFA, and ALT. The predicted gene content related to synthesis and degradation of ketone bodies, beta Alanine metabolism, glycosyltransferases, and other ion coupled transporters positively associated with BMI, body fat rate, and waistline, but negatively associated with body water rate, protein rate, muscle rate, and mineral salts rate.

**Figure 3.**
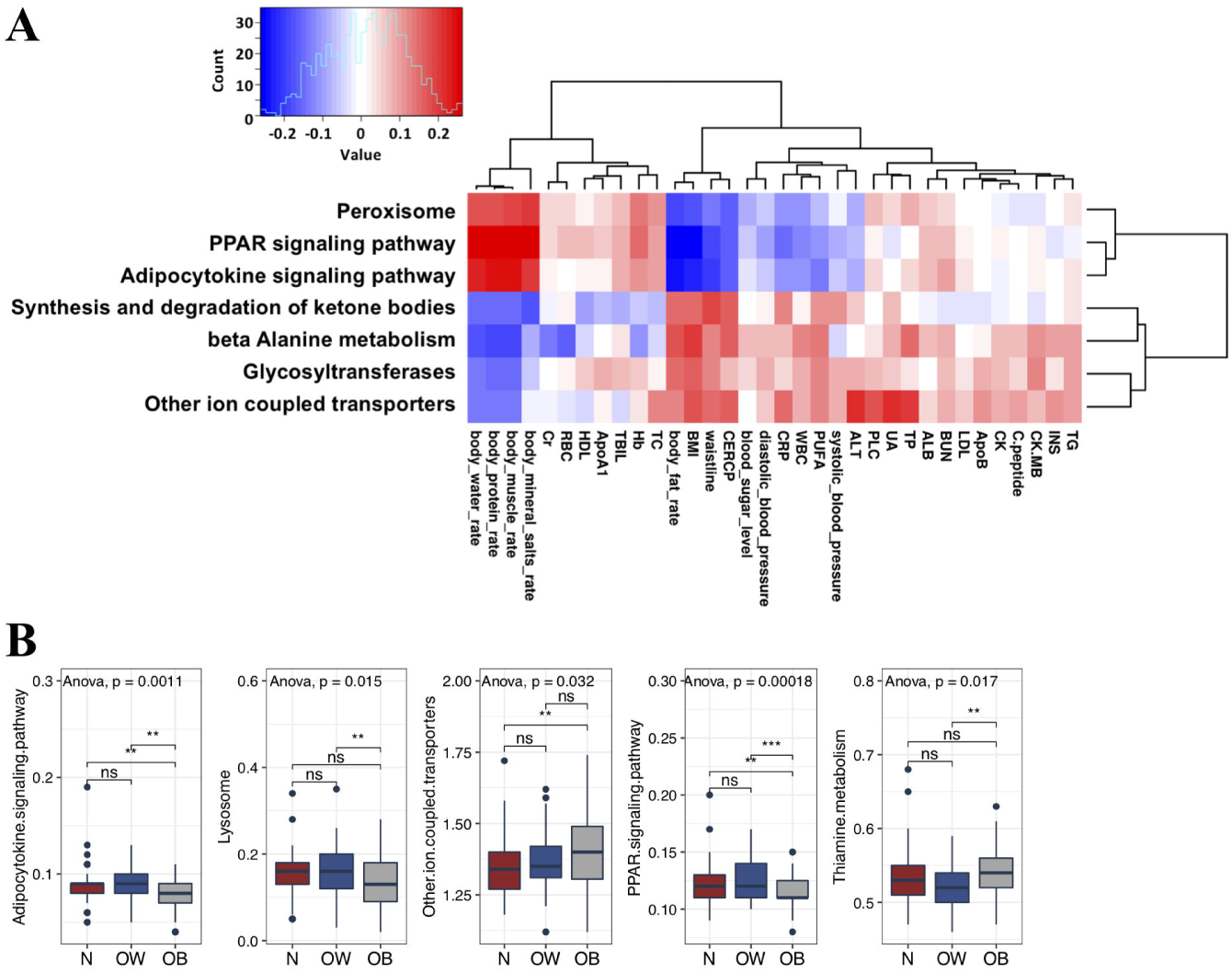
Functional divergence of gut microbiota across different BMI groups. **(A)** Predicted KEGG pathways that significantly correlated with BMI, and their associations with the boy composition and serum biochemical parameters. **(B)** KEGG pathways that are differentially expressed by the gut microbiome of the BMI categories. Each bar plot indicates the mean proportion of sequences assigned to a feature in each group. Whiskers represent 1.5* inter-quartile range. Relative abundances of were analyzed by one-way ANOVA followed by Wilcoxon test for pairwise comparisons. **P < 0.01 and ***P < 0.001. N, normal weight; OW, overweight; OB, obese.

### BMI Differences in The Gut Microbiome of Adolescents Are Influenced By Gender

With the gut microbiota sequencing data of 516 Chinese adult, we previously found that BMI differences in the gut microbiome composition are gender specific (Gao et al. 2018). Since age has also been reported as a confounding factor influencing the gut microbiome, and it is uncertain whether the gender differences discovered in adult people also exist in adolescents. Here we evaluate whether BMI differences in the gut microbiome of adolescents are influenced by gender. Neither alpha diversity nor beta diversity was found significantly different between Chinese boys and girls (Figure S1, S2). At the family level, the relative abundances of *Bacteroidaceae, Brucellaceae, Clostridiaceae, Lachnospiraceae, Planococcaceae*, and *Streptococcaceae* were more abundant in boys than girls (Figure S3A).

We then compared the relative abundances of the gut microbiome among the three BMI categories with gender stratification (Figure S3B). At the family level, a higher abundance of *Cytophagaceae* was observed in both obese girls and boys. Overweight boys had a lower abundance of *Actinomycetaceae*, and a higher abundance of *Nocardiaceae* than those with normal weight as well as obesity. At the genus level, both obese boys and girls had higher relative abundances of *Hymenobacter* and *Megasphaera*. A lower relative abundance of *Odoribacter* was particularly found in the gut microbiome of obese boys. In addition, *Actinomyces* and *Rhodococcus* were significantly lower and higher in overweight boys, respectively.

PICRUSt analysis revealed a higher level of bisphenol degradation in boys. A reduced level of tropane piperidine and pyridine alkaloid biosynthesis was only observed in obese boys (Figure S4). Obese girls had an increased level of selenocompound metabolism, and decreased levels of adipocytokine signaling pathway and ethylbenzene degradation. A lower level of PPAR signaling pathway was detected in both genders.

## DISCUSSION

The identification of gut microbial signatures that are responsible of obesity has great potential in prevention and treatment of overweight and obesity. Here we report BMI-associated patterns in the adolescent gut microbiome composition and functions. The imbalance of *Bacteroidetes* to *Firmicutes* ratio has been reported in numerous studies (Duncan et al, 2008; Jumpertz et al. 2011; Schwiertz et al. 2011), and we previously reported that Bacteroidetes was enriched in obese adults comparing to lean subjects (Gao et al. 2018). In the present data, no difference in the relative abundances of these two phyla upon comparison of obese and normal-weight adolescents, which is in agreement with findings of (Duncan et al. 2008; Jumpertz et al. 2011). We identified 56 microbial taxa significantly (p<0.01) correlated with BMI, and a large proportion of them belonging to Firmicutes phylum, especially families *Ruminococcaceae* and *Lachnospiraceae*. Supervised learning algorithm constructed based on the 56 BMI-associated OTUs resulted in the successful classification of obese adolescents with a high accuracy exceeding 90%.

The enrichment of *Ruminococcaceae* has been observed in animals fed by high-fat diet (Schwiertz et al. 2011), but some *Ruminococcaceae* taxa were also linked to a lower risk of weight gain (Menni et al. 2017). In human studies, higher abundances of *Ruminococcus* species (such as *Ruminococcus bromii* and *Ruminococcus obeum*) were also observed in obese subjects (Kasai et al. 2015), however, some *Ruminococcaceae* bacteria such as *Dialister, Methanobrevibacter*, and *Oscillospira* have also been associated with lower BMI (Jumpertz et al. 2011; Goodrich et al. 2014). From our data, we identified 17 OTUs assigned to Ruminococcaceae were significantly correlated with BMI, and many of which were inversely correlated with BMI including the well-studied butyrate producer *Faecalibacterium prausnitzii*, which is consistent with several studies (Borgo et al. 2018; Tilg and Moschen. 2014). Butyrate has been known exerting a profound effect in regulating metabolic inflammation, and reduced level of butyrate may contribute to low-grade chronic inflammation that participating the development of obesity. Nevertheless, there were also some *Ruminococcaceae* members positively correlated with BMI. Thus, investigations at higher phylogenetic level are needed to identify the specific bacteria species in terms of influence in weight gain.

Animal studies demonstrated that the abundance of family *Lachnospiraceae* increased along with body weight of mice fed with high-fat diet (Ravussin et al. 2012); colonization of *Lachnospiraceae* in germ-free mice induced significant increases in fasting blood glucose concentrations as well liver and mesenteric adipose weights, and reductions in plasma insulin levels. A human study with 190 Mexican children showed that the family *Lachnospiraceae* was significantly increased in overweight and obese children (Murugesan et al. 2015). Taken together, *Lachnospiraceae* may play a pivotal role in the development of obesity and type 2 diabetes (Kameyama et al. 2014). From our data, significant associations were obtained between 12 taxa of *Lachnospiraceae* and BMI. Similar to what we observed in *Ruminococcaceae*, both positive (8 OTUs) and negative (4 OTUs) correlations were detected between BMI and the *Lachnospiraceae* taxa. Hence, the specific species and strains are needed to be determined in terms of their influences in blood glucose level and weight gain.

Although the exact mechanisms by which the gut microbiota contributes to obesity are unclear, it is well established that modification of the gut microbiota can increase energy production, trigger low-grade inflammation, induce insulin resistance, and affect fatty acid tissue composition (Musso et al. 2011). Based on the inferred functional profiles of the fecal microbiome, we found that genes associated PPAR and adipocytokine signaling pathways were inversely correlated with CER/CP, blood sugar level, (both systolic and diastolic) blood pressure, CRP, WBC, PUFA, and ALT, and their levels were significantly downregulated in obese adolescents. PPARs are a member of the nuclear receptor superfamily of ligand-dependent transcription factors that is predominantly expressed in adipose tissue and the intestines, and it plays the master role in adipogenesis for obesity development (Lefterova et al. 2014). Lower levels of PPARs have been observed in obese patients, and activation of PPARs could decrease fibro-inflammation and ectopic fat accumulation in the adipose tissue (Corrales et al. 2018). Decline in bacterial genes capable of altering PPAR signaling may reflect a reduction of PPARs expression in the host. In fact, some gut bacteria have been found with capacities of regulating PPARs. For example, mice artificially infected with *Trichinella spiralis* could induce a decrease in the levels of PPARγ in the colon, which is accompanied by a decline in beneficial species (such as *Akkermansia*) and increase in pathogenic bacteria (such as *Escherichia*/*Shigella*) (Chen et al. 2017). Another pathway of importance that was differentially decreased in the obese adolescent was adipocytokine signaling. Adipocytokines derived from adipose tissue, including various hormones (such as leptin, adiponectin, resistin, visfatin) and cytokines (such as interleukin-6 and tumor necrosis factor α), are important regulators of energy homeostasis and mediators of inflammation and immunity (Tilg and Moschen. 2006). There is overwhelming evidence that deficiencies in adipocytokines contribute to the development of obesity and the associated comorbidities (Cao. 2014). Importantly, some adipocytokines are able to modulate gut microbial composition independently of dietary (Rajala et al. 2014). Further investigations are needed to identify the exact bacterial products that affect these two pathways in order to determine their roles in adiposity.

In our previous study with the fecal microbiome profiles Chinese adults (527 adults aged 37.3± 16.3), enrichment of *Fusobacteria* and *Actinobacteria* were observed in the male and female obese subjects, respectively (Gao et al. 2018). Functionally, bacterial genes associated with butyrate-acetoacetate CoA-transferase were found to be enriched in the gut microbiome of obese Chinese adults (Gao et al. 2018). However, these features were not obtained from the adolescent data (205 adolescents aged 13.31±1.55). These inconsistencies may relate to differences of age. The influence of gender on the gut microbiota have also been investigated in studies from ours and others. For example, members of *Bacteroides* were found in a lower level in adult females than males from surveys of European (Mueller et al. 2006) and US populations (Dominianni et al. 2015), which was also observed from our data of Shenzhen adolescents, but not in our previous study with Chinese adults (Gao et al. 2018). Indeed, geographical location has been found exerting a strong effect on human gut microbiota variations (He et al. 2018). Shenzhen is one of the most developed cities in China and world-wide, and its lifestyles (especially diet) are much similar to western countries than other areas of China. Hence, some similarities the gut microbiome composition may exist between Shenzhen citizens and westerners. Taken together, the influence of geographical location, age, and gender can (partly) explain the inconsistent patterns across studies.

In conclusion, the BMI-associated differences in gut microbiota profiles were able to be used as biomarkers for characterizing obese adolescents. The gene contents associated with PPAR signaling pathway significantly reduced in the gut microbiome of obese adolescents, and determining the exact metabolites produced by specific species may provide invaluable microbial targets for prevention, assessment, and treatment of obesity for adolescents. Our result reinforced a need to consider the influence of age, gender and geographical location when choosing controls to compare the gut microbiome of disease cases.

## DATA AVAILABILITY

Raw sequencing data have been deposited on the European Nucleotide Archive server, with accession number PRJEB33385.

## ACKNOWLEDGEMENT

We acknowledge the contributions of the nursing staff of Shenzhen Children’s Hospital for their help with blood, stool, and metadata collection. We thank all volunteers for their participation in this study.

## AUTHOR CONTRIBUTIONS

YL designed the study; BW, YP, MZ and SZ collected the samples; YP performed the experiment; XG and YC analyzed the data; XG, BW, YL and YC prepared the original draft; all authors read and approved the final version of the manuscript.

## COMPLIANCE WITH ETHICAL STANDARDS

### Funding

This study was funded by the Shenzhen key technology R&D program (JSGG20170413152936281, 20170502165510880), and University of Electronic Science and Technology of China (Y03019023601008022).

### Ethical approval

All procedures performed in studies involving human participants were in accordance with the ethical standards of the institutional and/or national research committee and with the 1964 Helsinki declaration and its later amendments or comparable ethical standards.

