## Supplementary material for "Gut Microbiome Biomarkers in Adolescent Obesity: a Regional Study"

Applied Microbiology and Biotechnology

**Supplementary Information**

**Table S1**. BMI-associated bacterial taxa are identified by Spearman correlations in all the subjects.

**Table S2.** BMI-associated bacterial taxa are identified by Spearman correlations in boys.

**Table S3.** BMI-associated bacterial taxa are identified by Spearman correlations in girls.

**Table S4.** BMI-associated PICRUSt-predicted KEGG pathways are identified by Spearman correlations in all the subjects.

**Table S5.** BMI-associated PICRUSt-predicted KEGG pathways are identified by Spearman correlations in boys.

**Table S6.** BMI-associated PICRUSt-predicted KEGG pathways are identified by Spearman correlations in girls.

**Figure S1**. Evaluating the gut microbiota biodiversity among the three BMI categories and for both gender groups. The **(A)** richness, **(B)** chao1 index, **(C)** Shannon index, and **(D)** Inverse Simpson index were measured at the OTU level. Comparisons were performed by one-way ANOVA followed by Wilcoxon test for pairwise comparisons. $*P < 0.05$, $**P < 0.01$, and $***P < 0.001$. N, normal weight; OW, overweight; OB, obese.

**Figure S2**. Evaluating effect of BMI and gender in explaining the variations in beta-dispersion. The compositional similarities between microbial communities (beta diversity) was estimated by PCoA based on Bray–Curtis distances of OTU relative abundances.

**Figure S3**. Differences in fecal microbiome composition are accessed with gender stratification. Bacterial taxa at the family and genus levels demonstrated difference in relative abundances of the gut microbiota **(A)** between boys and girls, and **(B)** across the three BMI groups with gender stratification. Each bar plot indicates the mean proportion of sequences assigned to a feature in each group. Whiskers represent 1.5* inter-quartile range. Relative abundances of were analyzed by one-way ANOVA followed by Wilcoxon test for pairwise comparisons. **P < 0.01 and ***P < 0.001. N, normal weight; OW, overweight; OB, obese.

**Figure S4**. Differences in fecal microbiome functions are accessed with gender stratification. Relative abundances of the PICRUSt-predicted pathways were compared **(A)** between boys and girls, and **(B)** across the three BMI groups with gender stratification. Each bar plot indicates the mean proportion of sequences assigned to a feature in each group. Whiskers represent 1.5* inter-quartile range. Relative abundances of were analyzed by one-way ANOVA followed by Wilcoxon pairwise comparisons. **P < 0.01 and ***P < 0.001. N, normal weight; OW, overweight; OB, obese.

**Table S1.** BMI-associated bacterial taxa are identified by Spearman correlations in all the subjects.

| Phylum | Family | OTU | R (Spearman index) | P-value | FDR | Adjusted P (Bonferroni) | Mean Abundance | Positive Samples |
|---|---|---|---|---|---|---|---|---|
| *Bacteroidetes* | *Bacteroidaceae* | p__*Bacteroidetes*__g__*Bacteroides*__s__*plebeius*_352617 | 0.26 | 1.29E-04 | 0.0056 | 0.065 | 1.74 | 118 |
| *Bacteroidetes* | *Bacteroidaceae* | p__*Bacteroidetes*__g__*Bacteroides*_213566 | 0.2 | 3.37E-03 | 0.042 | 1 | 1.04 | 76 |
| *Bacteroidetes* | *Bacteroidaceae* | p__*Bacteroidetes*__g__*Bacteroides*_313764 | -0.2 | 4.25E-03 | 0.047 | 1 | 1.41 | 166 |
| *Bacteroidetes* | *Bacteroidaceae* | p__*Bacteroidetes*__g__*Odoribacter*_4454586 | -0.18 | 9.07E-03 | 0.076 | 1 | 2.48 | 162 |
| *Firmicutes* | / | p__*Firmicutes*__o__*Clostridiales*_175650 | 0.2 | 4.19E-03 | 0.047 | 1 | 1.27 | 147 |
| *Firmicutes* | / | p__*Firmicutes*__o__*Clostridiales*_182167 | -0.33 | 9.65E-07 | 7.10E-05 | 0.00049 | 0.91 | 133 |
| *Firmicutes* | / | p__*Firmicutes*__o__*Clostridiales*_2230862 | 0.22 | 1.48E-03 | 0.027 | 0.73 | 0.66 | 73 |
| *Firmicutes* | / | p__*Firmicutes*__o__*Clostridiales*_358944 | -0.23 | 1.13E-03 | 0.022 | 0.56 | 0.85 | 85 |
| *Firmicutes* | / | p__*Firmicutes*__o__*Clostridiales*_359149 | 0.18 | 9.03E-03 | 0.076 | 1 | 1.13 | 147 |
| *Firmicutes* | / | p__*Firmicutes*__o__*Clostridiales*_470382 | 0.2 | 3.47E-03 | 0.042 | 1 | 1.19 | 167 |
| *Firmicutes* | / | p__*Firmicutes*__o__*Clostridiales*_553080 | -0.23 | 6.96E-04 | 0.017 | 0.35 | 1.02 | 141 |
| *Firmicutes* | *Clostridiaceae* | p__*Firmicutes*__f__*Clostridiaceae*_359750 | 0.23 | 1.12E-03 | 0.022 | 0.55 | 0.93 | 102 |
| *Firmicutes* | *Lachnospiraceae* | p__*Firmicutes*__f__*Lachnospiraceae*_208790 | 0.22 | 1.61E-03 | 0.028 | 0.79 | 1.97 | 131 |
| *Firmicutes* | *Lachnospiraceae* | p__*Firmicutes*__f__*Lachnospiraceae*_357312 | 0.25 | 2.59E-04 | 0.0089 | 0.13 | 2.13 | 172 |
| *Firmicutes* | *Lachnospiraceae* | p__*Firmicutes*__f__*Lachnospiraceae*_368698 | 0.22 | 1.73E-03 | 0.028 | 0.85 | 5.88 | 204 |
| *Firmicutes* | *Lachnospiraceae* | p__*Firmicutes*__f__*Lachnospiraceae*_514282 | -0.19 | 7.73E-03 | 0.07 | 1 | 2.88 | 198 |
| *Firmicutes* | *Lachnospiraceae* | p__*Firmicutes*__f__*Lachnospiraceae*_514996 | 0.25 | 2.32E-04 | 0.0086 | 0.12 | 5.29 | 196 |
| *Firmicutes* | *Lachnospiraceae* | p__*Firmicutes*__f__*Lachnospiraceae*_522107 | 0.45 | 1.23E-11 | 2.10E-09 | 6.30E-09 | 1.15 | 124 |
| *Firmicutes* | *Lachnospiraceae* | p__*Firmicutes*__f__*Lachnospiraceae*_542096 | -0.24 | 4.62E-04 | 0.013 | 0.23 | 1.7 | 179 |
| *Firmicutes* | *Lachnospiraceae* | p__*Firmicutes*__f__*Lachnospiraceae*_545473 | -0.26 | 2.15E-04 | 0.0086 | 0.11 | 2.02 | 183 |
| *Firmicutes* | *Lachnospiraceae* | p__*Firmicutes*__f__*Lachnospiraceae*_845273 | 0.21 | 2.43E-03 | 0.034 | 1 | 1.63 | 135 |

Continued

| Phylum | Family | OTU | R (Spearman index) | P-value | FDR | Adjusted P (Bonferroni) | Mean Abundance | Positive Samples |
|---|---|---|---|---|---|---|---|---|
| *Firmicutes* | *Lachnospiraceae* | p__*Firmicutes*__g__*Blautia*__s__*obeum*_514086 | -0.19 | 5.67E-03 | 0.059 | 1 | 0.87 | 145 |
| *Firmicutes* | *Lachnospiraceae* | p__*Firmicutes*__g__*Dorea*_363232 | 0.22 | 1.46E-03 | 0.027 | 0.72 | 0.82 | 122 |
| *Firmicutes* | *Lachnospiraceae* | p__*Firmicutes*__g__*Lachnospira*_320322 | 0.21 | 2.42E-03 | 0.034 | 1 | 0.89 | 119 |
| *Firmicutes* | *Mogibacteriaceae* | p__*Firmicutes*__f__*Mogibacteriaceae*_214036 | -0.2 | 3.52E-03 | 0.042 | 1 | 1.02 | 87 |
| *Firmicutes* | *Oscillospiraceae* | p__*Firmicutes*__g__*Oscillospira*_196831 | -0.18 | 8.45E-03 | 0.075 | 1 | 3.29 | 187 |
| *Firmicutes* | *Oscillospiraceae* | p__*Firmicutes*__g__*Oscillospira*_213885 | -0.34 | 5.02E-07 | 4.30E-05 | 0.00026 | 1.1 | 116 |
| *Firmicutes* | *Oscillospiraceae* | p__*Firmicutes*__g__*Oscillospira*_370046 | -0.2 | 3.26E-03 | 0.042 | 1 | 1.32 | 125 |
| *Firmicutes* | *Oscillospiraceae* | p__*Firmicutes*__g__*Oscillospira*_548032 | -0.25 | 2.99E-04 | 0.0097 | 0.15 | 1.35 | 122 |
| *Firmicutes* | *Ruminococcaceae* | p__*Firmicutes*__f__*Ruminococcaceae*_1062061 | -0.27 | 1.19E-04 | 0.0056 | 0.061 | 1.49 | 155 |
| *Firmicutes* | *Ruminococcaceae* | p__*Firmicutes*__f__*Ruminococcaceae*_180182 | -0.18 | 8.75E-03 | 0.076 | 1 | 1.14 | 160 |
| *Firmicutes* | *Ruminococcaceae* | p__*Firmicutes*__f__*Ruminococcaceae*_191928 | -0.21 | 2.95E-03 | 0.039 | 1 | 1.83 | 190 |
| *Firmicutes* | *Ruminococcaceae* | p__*Firmicutes*__f__*Ruminococcaceae*_193572 | 0.5 | 3.49E-14 | 9.00E-12 | 1.80E-11 | 0.96 | 94 |
| *Firmicutes* | *Ruminococcaceae* | p__*Firmicutes*__f__*Ruminococcaceae*_195436 | 0.54 | 5.81E-17 | 3.00E-14 | 3.00E-14 | 1.38 | 126 |
| *Firmicutes* | *Ruminococcaceae* | p__*Firmicutes*__f__*Ruminococcaceae*_195651 | -0.23 | 9.39E-04 | 0.02 | 0.46 | 1.09 | 167 |
| *Firmicutes* | *Ruminococcaceae* | p__*Firmicutes*__f__*Ruminococcaceae*_195757 | 0.32 | 3.73E-06 | 0.00021 | 0.0019 | 3.99 | 198 |
| *Firmicutes* | *Ruminococcaceae* | p__*Firmicutes*__f__*Ruminococcaceae*_199047 | -0.2 | 3.94E-03 | 0.046 | 1 | 2.22 | 181 |
| *Firmicutes* | *Ruminococcaceae* | p__*Firmicutes*__f__*Ruminococcaceae*_2066056 | -0.2 | 4.61E-03 | 0.05 | 1 | 0.89 | 107 |
| *Firmicutes* | *Ruminococcaceae* | p__*Firmicutes*__f__*Ruminococcaceae*_267718 | -0.23 | 9.03E-04 | 0.02 | 0.45 | 1.11 | 167 |
| *Firmicutes* | *Ruminococcaceae* | p__*Firmicutes*__f__*Ruminococcaceae*_362078 | -0.25 | 3.83E-04 | 0.012 | 0.19 | 4.97 | 193 |
| *Firmicutes* | *Ruminococcaceae* | p__*Firmicutes*__f__*Ruminococcaceae*_366352 | -0.22 | 1.85E-03 | 0.029 | 0.9 | 2.92 | 156 |
| *Firmicutes* | *Ruminococcaceae* | p__*Firmicutes*__f__*Ruminococcaceae*_558444 | -0.24 | 6.14E-04 | 0.016 | 0.31 | 0.69 | 100 |
| *Firmicutes* | *Ruminococcaceae* | p__*Firmicutes*__f__*Ruminococcaceae*_758482 | -0.23 | 8.72E-04 | 0.02 | 0.43 | 2.16 | 140 |

Continued

| Phylum | Family | OTU | R (Spearman index) | P-value | FDR | Adjusted P (Bonferroni) | Mean Abundance | Positive Samples |
|---|---|---|---|---|---|---|---|---|
| *Firmicutes* | *Ruminococcaceae* | p__*Firmicutes*__g__*Faecalibacterium*__s__*prausnitzii*_366068 | -0.18 | 8.94E-03 | 0.076 | 1 | 1.21 | 176 |
| *Firmicutes* | *Ruminococcaceae* | p__*Firmicutes*__g__*Faecalibacterium*__s__*prausnitzii*_525215 | -0.19 | 6.60E-03 | 0.063 | 1 | 4.63 | 204 |
| *Firmicutes* | *Ruminococcaceae* | p__*Firmicutes*__g__*Faecalibacterium*__s__*prausnitzii*_851865 | -0.19 | 5.91E-03 | 0.059 | 1 | 7.11 | 204 |
| *Firmicutes* | *Streptococcaceae* | p__*Firmicutes*__g__*Streptococcus*_579608 | 0.22 | 1.76E-03 | 0.028 | 0.85 | 4.91 | 205 |
| *Firmicutes* | *Streptococcaceae* | p__*Firmicutes*__g__*Streptococcus*_593803 | 0.19 | 5.81E-03 | 0.059 | 1 | 2.4 | 190 |
| *Firmicutes* | *Veillonellaceae* | p__*Firmicutes*__g__*Dialister*_583746 | 0.19 | 5.23E-03 | 0.055 | 1 | 0.65 | 73 |
| *Firmicutes* | *Veillonellaceae* | p__*Firmicutes*__g__*Megasphaera*_264967 | 0.29 | 1.90E-05 | 0.00099 | 0.0097 | 2.8 | 129 |
| *Firmicutes* | *Veillonellaceae* | p__*Firmicutes*__g__*Megasphaera*_266210 | 0.33 | 1.15E-06 | 7.40E-05 | 0.00059 | 3.18 | 143 |
| *Firmicutes* | *Veillonellaceae* | p__*Firmicutes*__g__*Megasphaera*_817140 | 0.23 | 1.13E-03 | 0.022 | 0.56 | 0.94 | 65 |
| *Fusobacteria* | *Fusobacteriaceae* | p__*Fusobacteria*__g__*Fusobacterium*_572889 | 0.21 | 2.04E-03 | 0.03 | 0.99 | 1.22 | 85 |
| *Proteobacteria* | *Enterobacteriacea* | p__*Proteobacteria*__f__*Enterobacteriaceae*_559204 | 0.21 | 2.05E-03 | 0.03 | 0.99 | 1.21 | 113 |
| *Proteobacteria* | *Enterobacteriaceae* | p__*Proteobacteria*__g__*Erwinia*_922761 | 0.2 | 4.31E-03 | 0.047 | 1 | 1.78 | 137 |
| *Proteobacteria* | *Enterobacteriaceae* | p__*Proteobacteria*__g__*Klebsiella*_813217 | 0.19 | 6.21E-03 | 0.061 | 1 | 0.92 | 83 |

**Table S2**. BMI-associated bacterial taxa are identified by Spearman correlations in boys.

| Phylum | Family | OTU | R (Spearman index) | P-value | FDR | Adjusted P (Bonferroni) | Mean Abundance | Positive Samples |
|---|---|---|---|---|---|---|---|---|
| *Actinobacteria* | *Bifidobacteriaceae* | p__*Actinobacteria*__g__*Bifidobacterium*__s__*bifidum*_365385 | 0.22 | 0.009244129 | 0.19 | 1 | 1.9 | 58 |
| *Bacteroidetes* | *Bacteroidaceae* | p__*Bacteroidetes*__g__*Bacteroides*__s__*plebeius*_352617 | 0.3 | 0.000377664 | 0.027 | 0.18 | 2.48 | 83 |
| *Bacteroidetes* | *Bacteroidaceae* | p__*Bacteroidetes*__g__*Bacteroides*_313764 | -0.23 | 0.007018048 | 0.19 | 1 | 2.89 | 116 |
| *Bacteroidetes* | *Flavobacteriaceae* | p__*Bacteroidetes*__g__*Hymenobacter*_593226 | 0.43 | 1.26E-07 | 1.60E-05 | 6.20E-05 | 0.89 | 28 |
| *Bacteroidetes* | *Flavobacteriaceae* | p__*Bacteroidetes*__g__*Hymenobacter*_798861 | 0.41 | 8.61E-07 | 8.50E-05 | 0.00042 | 0.88 | 25 |
| *Firmicutes* | / | p__*Firmicutes*__o__*Clostridiales*_2230862 | 0.24 | 0.004096946 | 0.13 | 1 | 1.32 | 52 |
| *Firmicutes* | / | p__*Firmicutes*__o__*Clostridiales*_175650 | 0.22 | 0.008769623 | 0.19 | 1 | 2.45 | 98 |
| *Firmicutes* | *Erysipelotrichaceae* | p__*Firmicutes*__f__*Erysipelotrichaceae*_145801 | -0.25 | 0.003650075 | 0.13 | 1 | 1.56 | 67 |
| *Firmicutes* | *Lachnospiraceae* | p__*Firmicutes*__f__*Lachnospiraceae*_522107 | 0.46 | 1.19E-08 | 2.00E-06 | 5.80E-06 | 2.44 | 87 |
| *Firmicutes* | *Lachnospiraceae* | p__*Firmicutes*__f__*Lachnospiraceae*_514996 | 0.3 | 0.000317367 | 0.026 | 0.15 | 7.53 | 132 |
| *Firmicutes* | *Lachnospiraceae* | p__*Firmicutes*__f__*Lachnospiraceae*_357312 | 0.27 | 0.001301343 | 0.064 | 0.63 | 3.7 | 113 |
| *Firmicutes* | *Lachnospiraceae* | p__*Firmicutes*__f__*Lachnospiraceae*_368698 | 0.23 | 0.006460895 | 0.19 | 1 | 8.01 | 137 |
| *Firmicutes* | *Lachnospiraceae* | p__*Firmicutes*__g__*Roseburia*_528362 | 0.22 | 0.008292749 | 0.19 | 1 | 3.56 | 111 |
| *Firmicutes* | *Lachnospiraceae* | p__*Firmicutes*__f__*Lachnospiraceae*_183804 | 0.22 | 0.008382515 | 0.19 | 1 | 2.8 | 108 |
| *Firmicutes* | *Lachnospiraceae* | p__*Firmicutes*__f__*Lachnospiraceae*_545473 | -0.22 | 0.009780073 | 0.19 | 1 | 3.38 | 121 |
| *Firmicutes* | *Mogibacteriaceae* | p__*Firmicutes*__f__*Mogibacteriaceae*_214036 | -0.22 | 0.008635773 | 0.19 | 1 | 1.58 | 53 |
| *Firmicutes* | *Oscillospiraceae* | p__*Firmicutes*__g__*Oscillospira*_213885 | -0.25 | 0.002949031 | 0.11 | 1 | 1.65 | 68 |
| *Firmicutes* | *Ruminococcaceae* | p__*Firmicutes*__f__*Ruminococcaceae*_195757 | 0.28 | 0.001005213 | 0.062 | 0.49 | 6.09 | 130 |
| *Firmicutes* | *Ruminococcaceae* | p__*Firmicutes*__f__*Ruminococcaceae*_195436 | 0.55 | 3.29E-12 | 1.30E-09 | 1.60E-09 | 2.8 | 91 |
| *Firmicutes* | *Ruminococcaceae* | p__*Firmicutes*__f__*Ruminococcaceae*_193572 | 0.55 | 5.32E-12 | 1.30E-09 | 2.60E-09 | 1.96 | 67 |
| *Firmicutes* | *Veillonellaceae* | p__*Firmicutes*__g__*Megasphaera*_266210 | 0.27 | 0.001177757 | 0.064 | 0.57 | 4.52 | 100 |
| *Firmicutes* | *Veillonellaceae* | p__*Firmicutes*__g__*Megasphaera*_264967 | 0.26 | 0.002477993 | 0.11 | 1 | 3.94 | 86 |

**Table S3**. BMI-associated bacterial taxa are identified by Spearman correlations in girls.

| Phylum | Family | OTU | R (Spearman index) | P-value | FDR | Adjusted P (Bonferroni) | Mean Abundance | Positive Samples |
|---|---|---|---|---|---|---|---|---|
| *Firmicutes* | / | p__*Firmicutes*__o__*Clostridiales*_470382 | 0.32 | 7.18E-03 | 0.18 | 1 | 2.28 | 58 |
| *Firmicutes* | / | p__*Firmicutes*__o__*Clostridiales*_182167 | -0.38 | 1.23E-03 | 0.093 | 0.64 | 2.61 | 53 |
| *Firmicutes* | *Clostridiaceae* | p__*Firmicutes*__f__*Clostridiaceae*_359750 | 0.4 | 7.46E-04 | 0.072 | 0.39 | 1.34 | 28 |
| *Firmicutes* | *Clostridiaceae* | p__*Firmicutes*__g__*SMB53*_555945 | 0.34 | 4.47E-03 | 0.14 | 1 | 5.7 | 67 |
| *Firmicutes* | *Lachnospiraceae* | p__*Firmicutes*__f__*Lachnospiraceae*_522107 | 0.36 | 2.49E-03 | 0.12 | 1 | 1.59 | 37 |
| *Firmicutes* | *Lachnospiraceae* | p__*Firmicutes*__f__*Lachnospiraceae*_542096 | -0.31 | 9.19E-03 | 0.19 | 1 | 3.43 | 64 |
| *Firmicutes* | *Ruminococcaceae* | p__*Firmicutes*__f__*Ruminococcaceae*_195436 | 0.46 | 8.24E-05 | 0.043 | 0.043 | 1.78 | 35 |
| *Firmicutes* | *Ruminococcaceae* | p__*Firmicutes*__f__*Ruminococcaceae*_193572 | 0.36 | 2.44E-03 | 0.12 | 1 | 1.33 | 27 |
| *Firmicutes* | *Ruminococcaceae* | p__*Firmicutes*__f__*Ruminococcaceae*_195757 | 0.35 | 3.03E-03 | 0.12 | 1 | 6.18 | 68 |
| *Firmicutes* | *Ruminococcaceae* | p__*Firmicutes*__g__*Oscillospira*_548032 | -0.32 | 7.26E-03 | 0.18 | 1 | 3.29 | 50 |
| *Firmicutes* | *Ruminococcaceae* | p__*Firmicutes*__f__*Ruminococcaceae*_187882 | -0.34 | 4.59E-03 | 0.14 | 1 | 1.39 | 29 |
| *Firmicutes* | *Ruminococcaceae* | p__*Firmicutes*__f__*Ruminococcaceae*_538322 | -0.34 | 4.13E-03 | 0.14 | 1 | 2.18 | 52 |
| *Firmicutes* | *Ruminococcaceae* | p__*Firmicutes*__f__*Ruminococcaceae*_212532 | -0.35 | 3.69E-03 | 0.14 | 1 | 2 | 40 |
| *Firmicutes* | *Ruminococcaceae* | p__*Firmicutes*__f__*Ruminococcaceae*_195651 | -0.37 | 1.85E-03 | 0.12 | 0.96 | 2.57 | 59 |
| *Firmicutes* | *Ruminococcaceae* | p__*Firmicutes*__f__*Ruminococcaceae*_366352 | -0.41 | 5.06E-04 | 0.066 | 0.26 | 4.5 | 50 |
| *Firmicutes* | *Ruminococcaceae* | p__*Firmicutes*__g__*Oscillospira*_213885 | -0.43 | 2.73E-04 | 0.066 | 0.14 | 2.97 | 48 |
| *Firmicutes* | *Streptococcaceae* | p__*Firmicutes*__g__*Streptococcus*_1078207 | 0.4 | 8.25E-04 | 0.072 | 0.43 | 2.66 | 60 |
| *Firmicutes* | *Veillonellaceae* | p__*Firmicutes*__g__*Megasphaera*_266210 | 0.42 | 3.80E-04 | 0.066 | 0.2 | 4.33 | 43 |
| *Firmicutes* | *Veillonellaceae* | p__*Firmicutes*__g__*Megasphaera*_264967 | 0.36 | 2.57E-03 | 0.12 | 1 | 3.9 | 43 |
| *Firmicutes* | *Veillonellaceae* | p__*Firmicutes*__g__*Megasphaera*_817140 | 0.32 | 7.65E-03 | 0.18 | 1 | 1.61 | 22 |
| *Fusobacteria* | *Fusobacteriaceae* | p__*Fusobacteria*__g__*Fusobacterium*_572889 | 0.31 | 9.21E-03 | 0.19 | 1 | 2.33 | 27 |
| *Proteobacteria* | *Enterobacteriaceae* | p__*Proteobacteria*__g__*Klebsiella*_813217 | 0.36 | 2.81E-03 | 0.12 | 1 | 1.55 | 29 |

Continued

| Phylum | Family | OTU | R (Spearman index) | P-value | FDR | Adjusted P (Bonferroni) | Mean Abundance | Positive Samples |
|---|---|---|---|---|---|---|---|---|
| *Proteobacteria* | *Enterobacteriaceae* | p__*Proteobacteria*__f__*Enterobacteriaceae*_1111294 | 0.33 | 5.52E-03 | 0.15 | 1 | 1.88 | 40 |
| *Proteobacteria* | *Enterobacteriaceae* | p__*Proteobacteria*__g__*Erwinia*_922761 | 0.32 | 8.76E-03 | 0.19 | 1 | 2.94 | 43 |

**Table S4**. BMI-associated PICRUSt-predicted KEGG pathways are identified by Spearman correlations in all the subjects.

| KEGG pathways | R (Spearman index) | P value | FDR | Mean Abundance | Positive Samples |
|---|---|---|---|---|---|
| PPAR signaling pathway | -0.31 | 6.72E-06 | 0.0014 | 3.9 | 206 |
| Adipocytokine signaling pathway | -0.23 | 8.00E-04 | 0.083 | 3.39 | 206 |
| Peroxisome | -0.22 | 1.62E-03 | 0.11 | 4.44 | 206 |
| Other ion coupled transporters | 0.2 | 4.33E-03 | 0.23 | 7.31 | 206 |
| Synthesis and degradation of ketone bodies | 0.19 | 6.23E-03 | 0.26 | 2.08 | 206 |
| beta Alanine metabolism | 0.18 | 8.30E-03 | 0.28 | 4.42 | 206 |
| Glycosyltransferases | 0.18 | 9.36E-03 | 0.28 | 5.34 | 206 |

**Table S5**. BMI-associated PICRUSt-predicted KEGG pathways are identified by Spearman correlations in boys.

| KEGG pathways | R (Spearman index) | P value | FDR | Mean Abundance | Positive Samples |
|---|---|---|---|---|---|
| PPAR signaling pathway | -0.3 | 3.53E-04 | 0.073 | 3.9 | 137 |
| Type I diabetes mellitus | -0.27 | 1.47E-03 | 0.15 | 2.87 | 137 |
| Valine leucine and isoleucine degradation | 0.24 | 4.84E-03 | 0.25 | 4.64 | 137 |
| Peroxisome | -0.23 | 5.86E-03 | 0.25 | 4.44 | 137 |
| Synthesis and degradation of ketone bodies | 0.23 | 6.45E-03 | 0.25 | 2.07 | 137 |
| Biosynthesis of vancomycin group antibiotics | -0.22 | 9.26E-03 | 0.25 | 3.15 | 137 |
| Adipocytokine signaling pathway | -0.22 | 9.82E-03 | 0.25 | 3.39 | 137 |

**Table S6**. BMI-associated PICRUSt-predicted KEGG pathways are identified by Spearman correlations in girls.

| KEGG pathways | R.(Spearman index) | P value | FDR | Mean Abundance | Positive Samples |
|---|---|---|---|---|---|
| PPAR signaling pathway | -0.35 | 3.61E-03 | 0.75 | 3.87 | 69 |

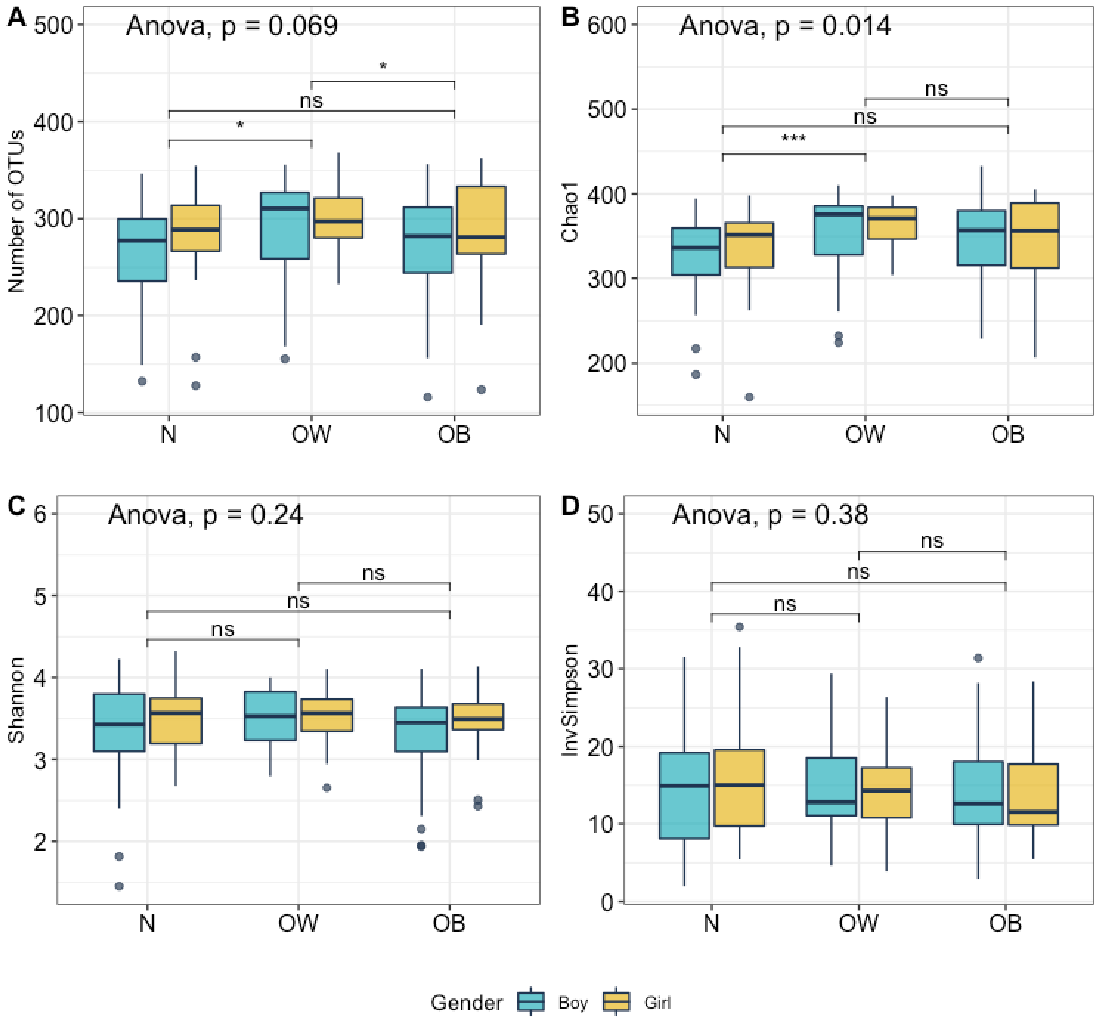


**Figure S1**. Evaluating the gut microbiota biodiversity among the three BMI categories and for both gender groups. The **(A)** richness, **(B)** chao1 index, **(C)** Shannon index, and **(D)** Inverse Simpson index were measured at the OTU level. Comparisons were performed by one-way ANOVA followed by Wilcoxon test for pairwise comparisons. *P < 0.05, **P < 0.01, and ***P < 0.001. N, normal weight; OW, overweight; OB, obese.

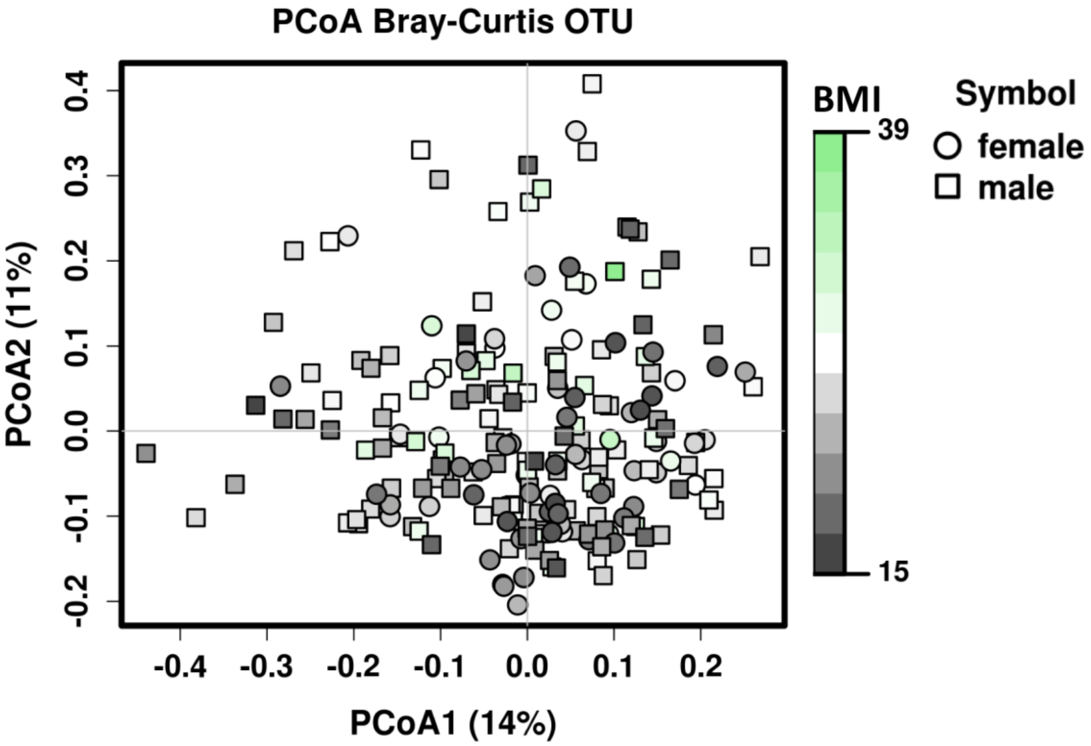


**Figure S2**. Evaluating effect of BMI and gender in explaining the variations in beta-dispersion. The compositional similarities between microbial communities (beta diversity) was estimated by PCoA based on Bray–Curtis distances of OTU relative abundances.

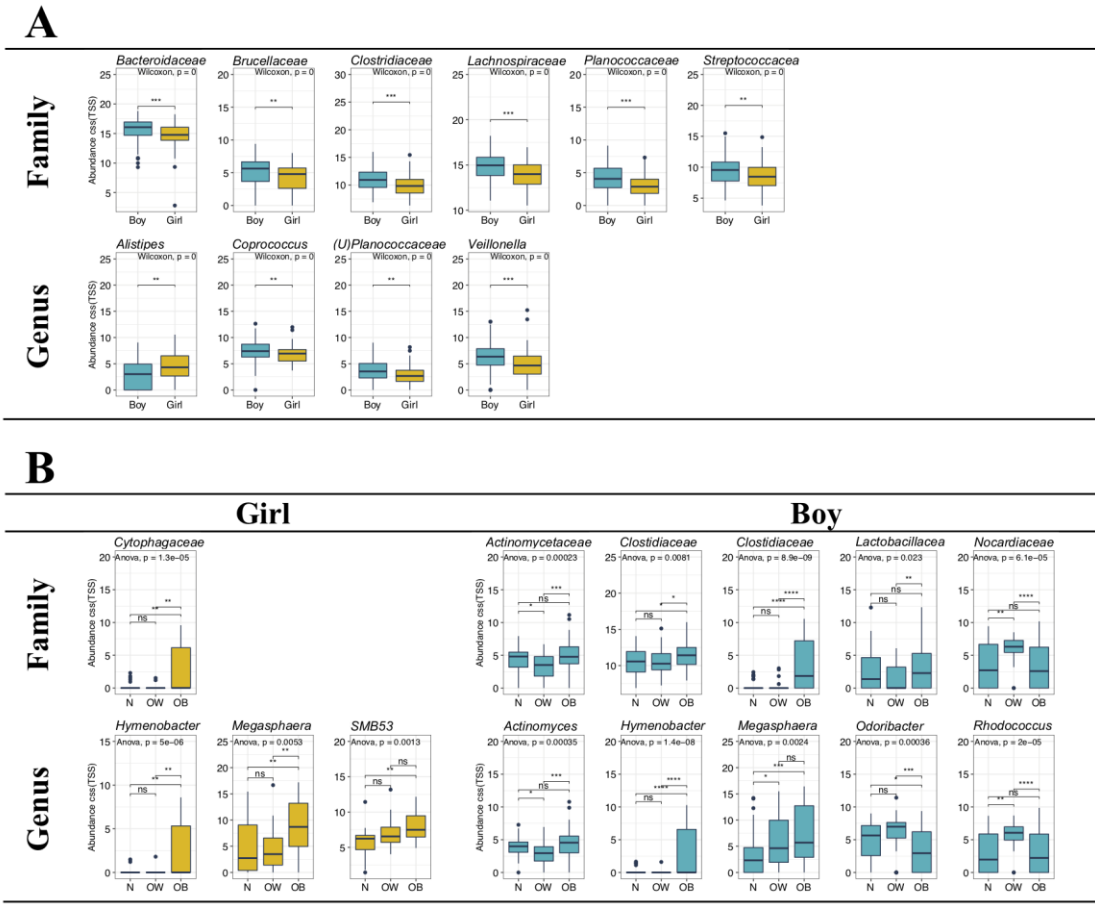


**Figure S3**. Differences in fecal microbiome composition are accessed with gender stratification. Bacterial taxa at the family and genus levels demonstrated difference in relative abundances of the gut microbiota **(A)** between boys and girls, and **(B)** across the three BMI groups with gender stratification. Each bar plot indicates the mean proportion of sequences assigned to a feature in each group. Whiskers represent 1.5* inter-quartile range. Relative abundances of were analyzed by one-way ANOVA followed by Wilcoxon test for pairwise comparisons. **P < 0.01 and ***P < 0.001. N, normal weight; OW, overweight; OB, obese.

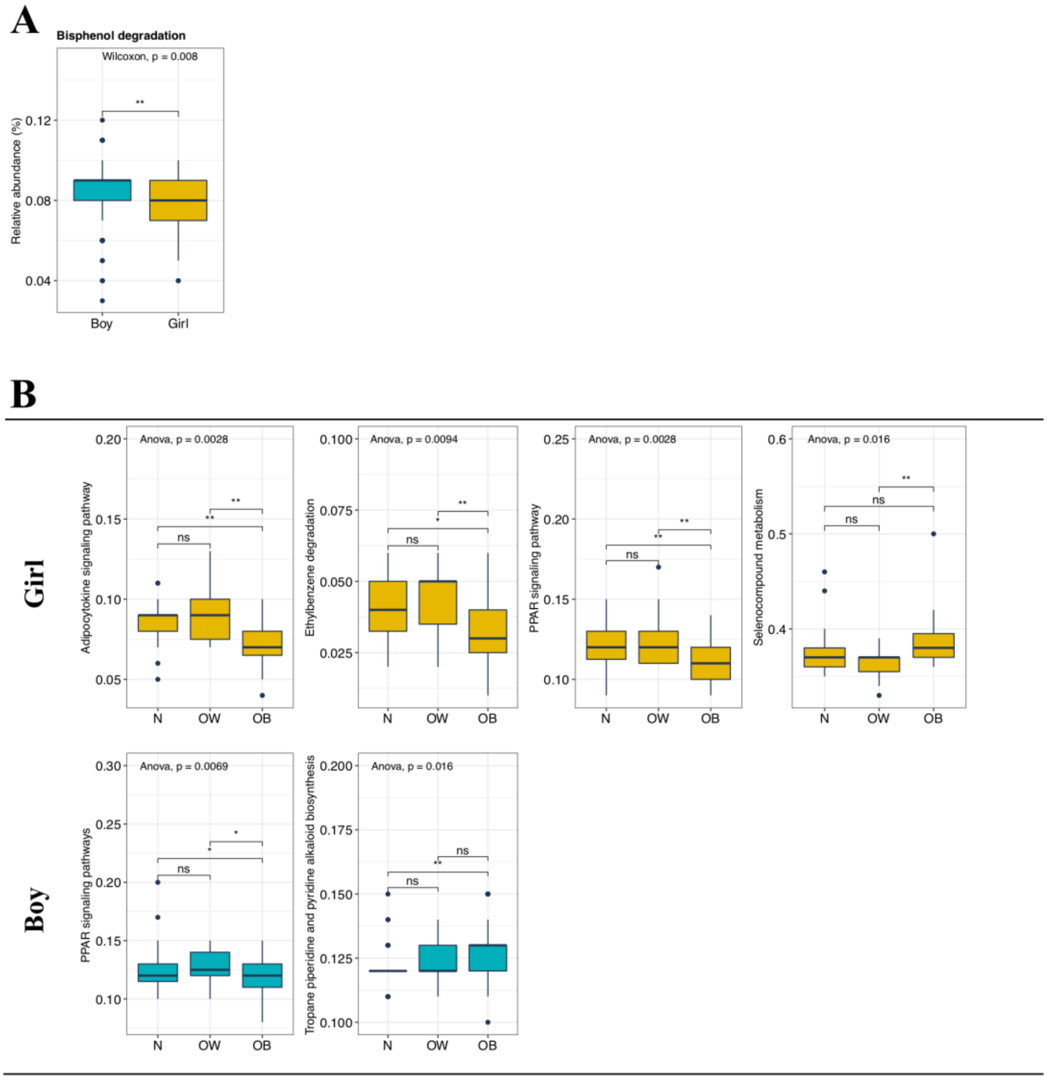


**Figure S4**. Differences in fecal microbiome functions are accessed with gender stratification. Relative abundances of the PICRUSt-predicted pathways were compared **(A)** between boys and girls, and **(B)** across the three BMI groups with gender stratification. Each bar plot indicates the mean proportion of sequences assigned to a feature in each group. Whiskers represent 1.5* inter-quartile range. Relative abundances of were analyzed by one-way ANOVA followed by Wilcoxon pairwise comparisons. **P < 0.01 and ***P < 0.001. N, normal weight; OW, overweight; OB, obese.
